# FIND: a software tool for identifying population-enriched pathogenic variants in gnomAD

**DOI:** 10.64898/2026.06.05.730273

**Authors:** Aaron L. Horowitz, Adam Z. Liebman, Susan W. Liebman

## Abstract

Identification of pathogenic population-enriched or founder mutations has facilitated efficient targeted screening. Founder mutations are a well-characterized subset of these, arising in a single ancestor and becoming enriched in a descendant population through bottlenecks and endogamy. However, many population-enriched variants that may or may not be founders also reveal population-specific disease burden and remain hidden in plain sight within existing datasets. To address this gap, we developed FIND (Founder-like candidates hidden IN Data), a web tool that identifies variants in gnomAD with frequencies >0.00008 in one ancestry group and at least tenfold higher than in all others (after zeroing populations with fewer than five observed alleles). The search is restricted to looking at pathogenic, likely pathogenic, and predicted loss-of-function variants. Testing FIND on the genes *FLNC, TMEM127, MYH7*, *BRCA1,* and *BRCA2* confirmed its utility and functionality by identifying twelve well-known founder or population-enriched mutations. In addition, FIND identified five variants previously reported as recurrent in a population but never compared across populations, and three never previously reported in any specific population. The population enrichment of all 20 variants was statistically significant, and a permutation test showed that FIND rarely returns results by chance. Variants enriched in African American and Admixed American populations were validated with the All of Us database, highlighting the utility of this approach for populations historically underrepresented in genetic studies. The source code is freely available at https://github.com/aacoder105/FIND under an MIT license, with a web interface at https://ethnic-variant-mutation-finder.onrender.com/.

## Introduction

Many diseases vary in both frequency and severity across populations [1]. Part of these differences are caused by the presence of mutations prevalent in a specific ancestry group and not in others. Some of these are pathogenic founder mutations that originated from a single or a few ancestors before becoming enriched in the descendant population. This status is generally confirmed by haplotype evidence [2] to distinguish them from variants that are simply population-enriched, i.e. present at higher frequency in one group than in others but with no evidence yet as to why.

Identifying population-enriched variants is useful because they can inform diagnostic decisions [3]. Knowing that a variant is ancestry-enriched implies that the mutation is not de novo and that others in the family are at increased risk for the disease, even without a known family history. Knowledge of population-enriched mutations can also alert physicians and patients to include the relevant gene in screening for patients of that ancestry. Founder-aware screening is now established clinical practice across several populations and diseases: in the Ashkenazi Jewish population, testing for three *BRCA1/BRCA2* founder variants is recommended for breast and ovarian cancer patients and accounts for the majority of identified pathogenic alleles in that group [4]; in the French Canadian population of Quebec, founder-variant panels are used as a first-line, cost-effective test [5]; and the Finnish disease heritage comprises roughly three dozen conditions, each driven largely by a single founder mutation, supporting healthcare-embedded carrier screening [6]. These precedents show the clinical value of founder-aware, population-specific screening, but they are largely limited to a small number of well-studied populations.

Other ancestry groups (e.g., African American) have not benefited to the same extent from population-specific screening, in part because fewer population-enriched variants have been identified in them, as less genetic testing and sequencing has been conducted in these groups [7]. FIND may help extend the benefit to these groups.

No general tool exists to mine gnomAD for population-enriched variants. Although recent studies have examined ancestry-enriched variants in specific gene sets, they have relied on bespoke analytic pipelines requiring programming expertise and, in some cases, credentialed biobank access [8]. We have developed and tested FIND, a Python tool that automates the tedious task of systematically screening gnomAD for population-enriched variants. FIND screens gnomAD pathogenic or likely pathogenic variants in any human gene, including known disease genes, for those that are population-enriched. In addition to pathogenic variants, FIND also flags predicted loss-of-function (pLoF) variants that are population-enriched, because even if not yet classified as pathogenic, they are plausible candidates for pathogenicity pending reclassification. With FIND, researchers and bench scientists do not need programming skills to uncover population-enriched pathogenic variants in genes of interest and export the results to a Microsoft Excel (.xlsx) file, helping to democratize use of the gnomAD database.

One challenge in working with ancestry groupings in population databases is that they are discrete approximations of continuous genetic ancestry. Their resolution depends on how well each population is represented, and recently admixed groups can mask substantial ancestry-specific variation [9]. The populations examined here are limited to the groups and data reported in gnomAD [1, 10].

## Materials and methods

FIND uses data from gnomAD v4.1 to look for population-enriched variants that have a reasonable chance of causing disease by focusing on variants currently categorized as pathogenic/likely pathogenic (P/LP) or loss of function (stop gained, frameshift, splice acceptor, splice donor on the gene’s canonical transcript) [11–14]. The pathogenicity categorization is taken from the ClinVar annotations provided within gnomAD, so FIND misses variants that cause disease but are not yet classified as P/LP in ClinVar. This is especially relevant for understudied populations where variant of unknown significance (VUS) classification lags the most. We chose three thresholds to balance sensitivity for population enriched variants against false positives from sampling noise. All three are adjustable parameters in the source code.

First, to reduce noise from sparsely sampled rare variants, FIND requires a minimum allele count of five, treating those below five as zero in both comparison and enriched populations. This threshold reduces sensitivity to low-count founders in under-sampled populations. Second, FIND requires enriched variants be at least tenfold more common in one ancestry group (Admixed American [Latino] AMR, African/African American AFR, Amish, Ashkenazi Jewish ASJ, East Asian EAS, Finnish FIN, Middle Eastern, Non-Finnish European NFE, South Asian) than in any other. This deliberately stringent threshold of tenfold captures pronounced enrichment rather than borderline differences. gnomAD’s tenth group, “Remaining,” which contains individuals not assigned to any of these nine ancestry groups, is excluded throughout, because it is not a single ancestry group and could mask or mimic enrichment. Third, FIND requires that the frequency in the enriched population exceed a minimum threshold. This value was set to 0.00008 to identify less common population-enriched dominant variants since most pathogenic variants have frequencies of <0.0001 [15]. Variants obtained using these thresholds are presented in Results. All three thresholds were set during development using *FLNC*, *TMEM127*, *MYH7* and *BRCA2*, as reported in our first preprint [16]. *BRCA1* was analyzed afterward with the thresholds unchanged and therefore serves as an independent test. When every comparison group falls below five alleles and is set to zero, the tenfold criterion is met automatically, and selection rests on the allele-count and frequency thresholds. The tenfold criterion is therefore a screening setting, not a test of statistical significance.

We assessed statistical support for each variant in Table 1 with a one-sided Fisher’s exact test comparing the allele count in the enriched group with the pooled count in the other eight groups (excluding the Remaining group). We used a one-sided test because our hypothesis is directional. We ask whether a variant is enriched in one group, not simply whether its frequency differs. These tests used raw gnomAD counts, not the zeroed counts FIND screens on. Because these 20 variants were selected from the 2,057 screened, we applied a Bonferroni correction (p < 0.05/2,057, or 2.4 × 10^-5^). Raw counts, fold ratios and p-values for each variant are given in S1 Table.

**Table 1.** Population-enriched pathogenic variants identified by FIND in five actionable genes (*FLNC, TMEM127, MYH7, BRCA*1 and *BRCA2*). For each variant, the table lists the gene; the ClinVar Variant ID and predicted protein change; the gnomAD v4.1 variant identifier and DNA change; the allele frequency in the enriched population (with the gnomAD ancestry group in parentheses); the category (defined below); the allele count and allele number in All of Us (NA where not checked); the ClinVar review status (0 to 4 stars, checked October 16, 2026); and references. Variants are separated into four categories: 1) previously established founders, reported in the literature on evidence beyond frequency, *Known Founder;* 2) previously shown enriched relative to other populations on frequency evidence, *Known Population-Enriched;* 3) previously reported as recurrent within a population but not compared across populations, *First cross-population comparison*; 4) not previously reported in any specific population, *First population-specific report*. Categories 1 and 2 confirm that FIND is working. Categories 3 and 4 are the novel findings; category 4 is the most novel. Frequencies are from gnomAD v4.1 after zeroing populations with four or fewer observed alleles. HGVS DNA descriptions reference the following transcripts: NM_001458.5 (*FLNC*), NM_017849.4 (*TMEM127*), NM_000257.4 (*MYH7*), NM_007294.4 (*BRCA1*), NM_000059.4 (*BRCA2*). All of Us was used to corroborate gnomAD enrichment for the African/African American and Admixed American groups, which All of Us oversamples; it was not used for East Asian, Ashkenazi Jewish, or Finnish variants (see Methods). Population abbreviations: AFR, African/African American; AMR, Admixed American (Latino); EAS, East Asian; NFE, Non-Finnish European; FIN, Finnish; ASJ, Ashkenazi Jewish. AC, allele count; AN, allele number. Raw allele counts and allele numbers for every gnomAD group, fold ratios, homozygote counts, gnomAD filter status and Fisher’s exact test p-values for each variant are given in S1 Table.

| Gene | ClinVar ID / Protein | gnomAD ID / DNA | Freq (enriched pop) | Category | All of Us (AC/AN) | ClinVar stars | Refs |
|---|---|---|---|---|---|---|---|
| *FLNC* | 420146/p.? | 7-128845989-G-C / c.3791-1G>C | 0.0001689 (ASJ) | Known Population-Enriched | NA | 2 | [21-23] |
| *TMEM127* | 571215/p.? | 2-96254998-C-G / c.245-1G>C | 0.00009341 (AFR) | First population-specific report | 10/189,216 AFR 10/1,071,394 total | 2 |  |
| *MYH7* | 155814/p.Arg1053Gln | 14-23422267-C-T / c.3158G>A | 0.0004372 (FIN) | Known Founder | NA | 3 | [24] |
| *MYH7* | 177667/p.Arg869His | 14-23424842-C-T / c.2606G>A | 0.0001333 (AFR) | First cross-population comparison | 28/189,214 AFR 40/1,071,280 total | 3 | [25] In Tuscany: population-enrichment not reported |
| *MYH7* | 14124/p.Glu497Asp | 14-23428587-C-A / c.1491G>T | 0.000236 (ASJ) | First population-specific report | NA | 2 |  |
| *BRCA1* | 17677/ p.Gln1756fs | 17-43057062-T-TG/ c.5266dup | 0.001183 (ASJ) | Known Founder | NA | 3 | [26] |
| *BRCA1* | 17662/p.Glu23fs | 17-43124027-ACT-A/ c.68_69del | 0.004158 (ASJ) | Known Founder | NA | 3 | [26] |
| *BRCA1* | 54381/p.Leu63Ter | 17-43106480-A-T/c.188T>A | 0.0001117 (EAS) | Known Founder | NA | 3 | [26, 27] |
| *BRCA1* | 54552/p.Glu797fs | 17-43093140-TTC-T/c.2389_2390del | 0.0001114 (EAS) | First cross-population comparison | NA | 3 | [27] In Japanese; population-enrichment not reported |
| *BRCA2* | 9326/p.Asn257LysfsTer17 | 13-32331003-ACAAAT-A / c.771_775del | 0.0001250 (FIN) | Known Founder | NA | 3 | [26, 28, 29] |
| *BRCA2* | 37830/p.Gln1089SerfsTer10 | 13-32337618-C-CT / c.3264dup | 0.0001260 (AMR) | First cross-population comparison | 43/194,852 AMR 44/1,071,280 Total | 3 | [30, 31] In Hispanic BRCA2 carriers; population-enrichment not reported |
| *BRCA2* | 38221/p.Val3079PhefsTer4 | 13-32380123-CG-C / c.9235del | 0.0001836 (AMR) | First cross-population comparison | 3/194,834 AMR 3/1,071,116 Total | 3 | [32, 33] In Peru, and Latin America, population-enrichment not reported |
| *BRCA2* | 9325/p.Ser1982ArgfsTer22 | 13-32340300-GT-G / c.5946del | 0.005339 (ASJ) | Known Founder | NA | 3 | [26, 34] |
| *BRCA2* | 37867/p.Glu1308Ter | 13-32338277-G-T / c.3922G>T | 0.0001779 (AMR) | Known Founder | 42/194,810 AMR 44/1,071,100 total | 3 | [35] |
| *BRCA2* | 37975/p.Ile1859LysfsTer3 | 13-32339928-CAATT-C / c.5576_5579del | 0.0001563 (EAS) | Known Founder | NA | 3 | [26, 36] |
| *BRCA2* | 9318/p.Leu2092ProfsTer7 | 13-32340629-CTT-C / c.6275_6276del | 0.00008747 (NFE) | Known Founder | NA | 3 | [26, 37] |
| *BRCA2* | 38076/p.Arg2318Ter | 13-32346841-C-T / c.6952C>T | 0.0003136 (EAS) | First cross-population comparison | NA | 3 | [38] In 3 Japanese; population-enrichment not reported. |
| *BRCA2* | 38099/p.Arg2494Ter | 13-32356472-C-T / c.7480C>T | 0.0001562 (FIN) | Known Founder | NA | 3 | [26, 29] |
| *BRCA2* | 52759/p.? | 13-32380005-A-G / c.9118-2A>G | 0.0002343 (FIN) | Known Founder | NA | 3 | [29] |
| *BRCA2* | 38260/p.Cys3233TrpfsTer15 | 13-32398209-TTGTA-T / c.9699_9702del | 0.000301 (AMR) | First population-specific report | 27/194,832 AMR 40/1,071,180 total | 3 |  |

These threshold choices reflect a deliberate tradeoff. We examined it with a sweep, varying the minimum allele count (3, 4, 6, 8 and 10 alleles) and the fold enrichment (5-, 15-, 20-, and 50-fold) one at a time while holding the other thresholds at their defaults (S2 Table). We queried gnomAD, ClinVar, and the All of Us public Data Browsers repeatedly, both manually and via FIND, between October 25, 2025 and June 21, 2026, as we collected data and refined classification thresholds, and again on October 15 and 16, 2026 for the analyses in Table 1 and S1–S3 Tables. All data used were publicly available in aggregate form; no individual-level or identifying participant information was accessed at any point.

To estimate how often FIND’s criteria return a variant by chance, we performed a permutation test. For each of the 2,057 screened variants, we kept the total allele count and randomly reassigned its alleles to the nine gnomAD groups in proportion to each group’s allele number for that variant, then applied FIND’s criteria. We repeated this 1,000 times and recorded results (S3 Table). The Fisher’s exact tests, threshold sweep and permutation test were run with Python scripts that apply FIND’s criteria to gnomAD browser exports downloaded on October 15, 2026; before use, the scripts were required to reproduce FIND’s own output for the five genes.

Up to ten genes can be searched together, and the returned variants can be exported to an Excel file. Fig 1 is a conceptual overview of FIND. In brief, FIND accepts up to ten gene symbols, retrieves each gene’s variants from gnomAD v4.1 through its GraphQL API, keeps variants that are pathogenic or likely pathogenic in ClinVar or predicted loss-of-function, computes ancestry-specific allele frequencies with counts below five set to zero, and returns variants meeting the enrichment and frequency criteria. A sample screenshot of FIND’s results for *BRCA2* (MIM:600185) [17] is shown in Fig 2.

**Fig 1.**
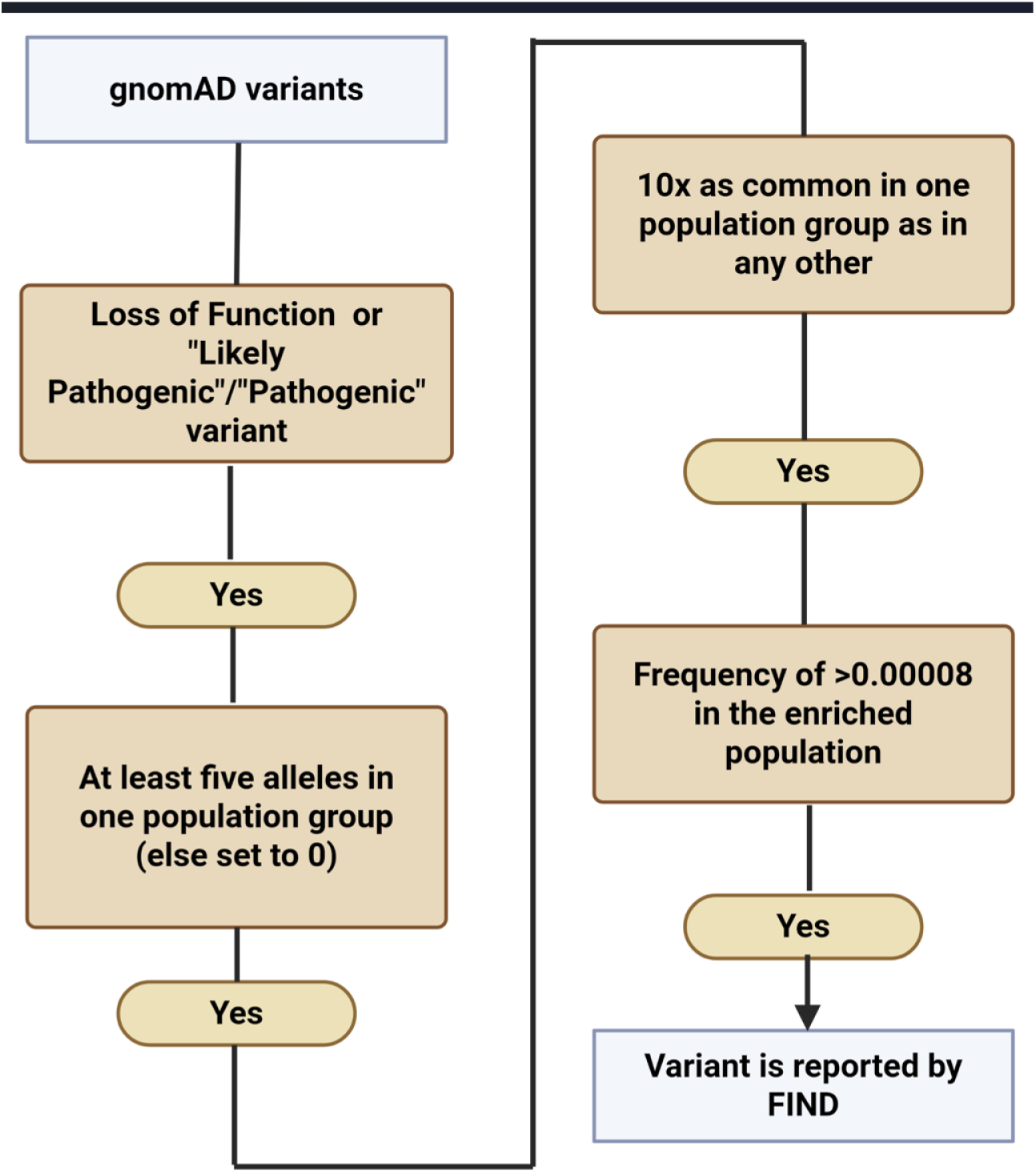
Visual representation of the logic of FIND. FIND retrieves variants of a queried gene from gnomAD v4.1 (or future v4 updates), choosing only those classified as pathogenic or likely pathogenic in ClinVar or predicted to be loss-of-function (stop-gained, frameshift, splice acceptor, splice donor). If the count of any variant in a population group is four or fewer, it is set to zero. Then, variants whose frequency in one population exceeds that in all others by at least tenfold while exceeding the 0.00008 minimum frequency threshold are selected. *Created in BioRender. Liebman, S. (2026)* https://BioRender.com/u2f39sg

**Fig 2.**
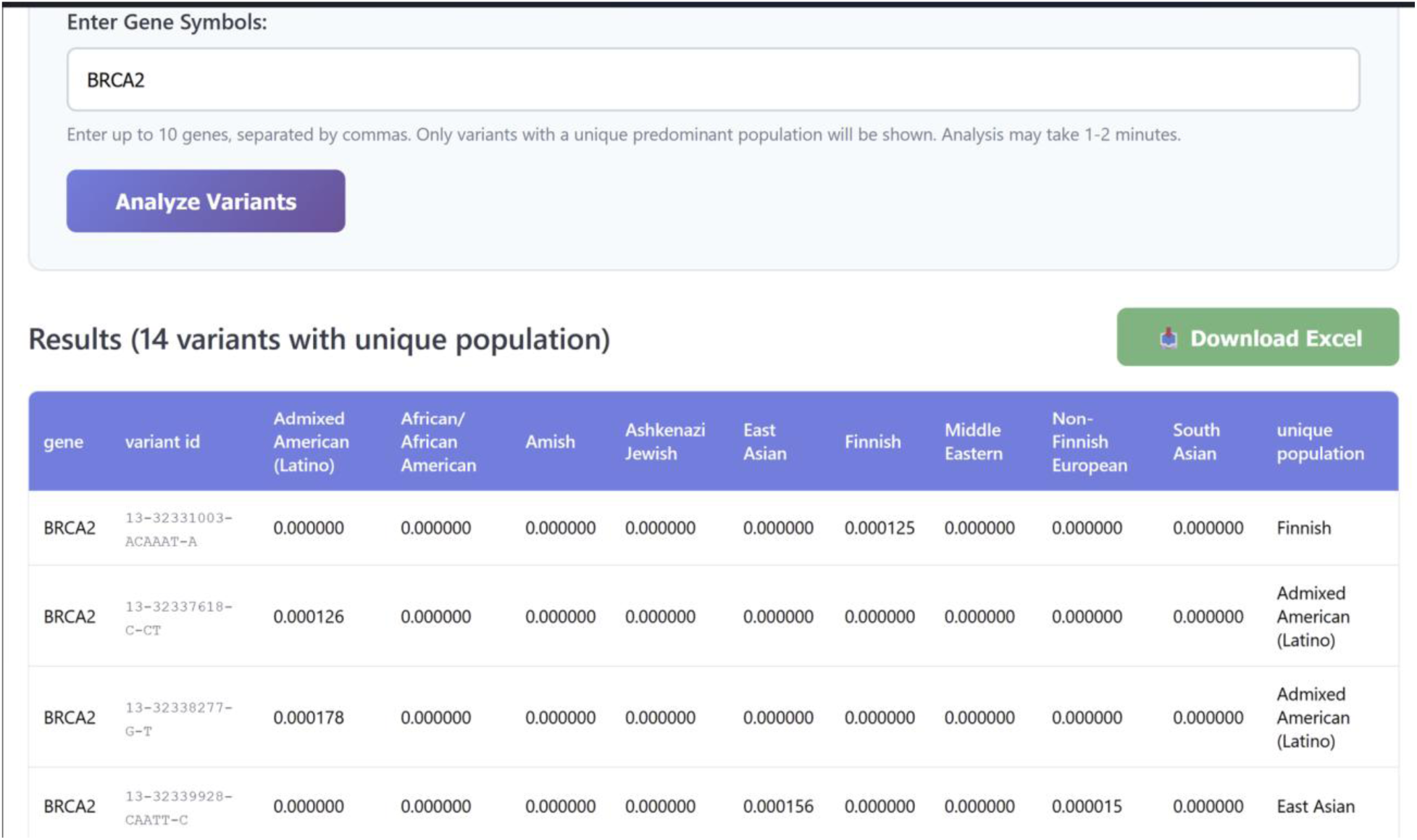
Screenshot of sample FIND results. Shown is a screen shot from a FIND query of the *BRCA2* gene. When on the web, more results can be viewed by scrolling. Up to 10 genes can be queried at once. The results table lists each variant by its gnomAD identifier and gives its allele frequency in each ancestry group (showing values of 0.000000 for populations where the variant was either absent or zeroed under the four-or-fewer-allele threshold). The last column identifies the unique ancestry group in which the variant is enriched (labeled ’unique population’ in the current interface). Results can be downloaded as an Excel file.

We manually corroborated whether the variants enriched in African/African American and Admixed American (Latino) populations were also enriched in All of Us [18]. gnomAD aggregates data from multiple research studies worldwide, totaling over 800,000 sequences, and is not designed to oversample any particular underrepresented population [11], whereas All of Us recruits only within the U.S. and deliberately oversamples populations historically underrepresented in biomedical research. The latest All of Us release provides whole-genome sequence data from approximately 414,000 individuals [19] including oversampling of the African/African American and Admixed American (Latino) populations, which is why we used it to corroborate enrichment in those groups. All of Us was not used for East Asian variants as known East Asian founder variants established in the literature are absent from All of Us despite roughly 25,000 East Asian participants, likely because All of Us samples US residents and under-represents the source populations in which these founders are concentrated. Ashkenazi Jewish and Finnish are not reported as separate ancestry groups in All of Us.

We tested FIND by searching for population-enriched pathogenic variants in five actionable genes [20]: *FLNC, BRCA2, BRCA1* were chosen because known founders or population enriched variants exist in these genes (validation true-positives); *MYH7* and *TMEM127* were chosen as additional actionable genes to test the tool more broadly. We didn’t continue to investigate loss-of-function variants that are not currently classified as pathogenic/likely pathogenic in ClinVar. Unless otherwise mentioned, each reported variant passed gnomAD v4.1 quality control with no observed homozygotes (PASS filter, clean allele balance and depth, no problematic site flags). Two reported variants (*BRCA2* p.Arg2318Ter and *BRCA1* p.Glu797ThrfsTer3) were observed in gnomAD’s exome dataset but not its (smaller) genome dataset. This reflects the larger size and different ancestry composition of the exome dataset rather than a quality concern.

We also considered a version of the software that applied the tenfold-enrichment criterion before the count-to-zero threshold, reversing the order used in our primary approach. This logic recovered nothing the primary approach had missed and failed to detect several variants the primary approach found: three *BRCA2* founders, one *MYH7* (MIM:160760) founder, and one *FLNC* (MIM:102565) known population-enriched variant, plus two *BRCA2* and one *MYH*7 among the previously unreported population-enriched variants. We therefore retained the original filter order.

Table 1 is based on a FIND search run on October 16, 2026, using gnomAD v4.1 and the ClinVar annotations it provides (ClinVar’s October 11, 2026 release). FIND’s raw output for the five genes contained 26 variants. Table 1 was produced from it in five steps: (1) pLoF variants classified as VUS or with conflicting classifications were excluded (two in *MYH7* and three in *BRCA2*; see Results); (2) one *BRCA1* variant failing gnomAD’s AS-VQSR filter was excluded; (3) gnomAD quality flags were reviewed for the remaining variants; (4) African/African American and Admixed American variants were checked in All of Us; and (5) each variant was assigned a category from the literature. Users should review FIND’s output in the same way. User output will include pLoF variants of uncertain significance and variants that fail gnomAD quality filters, and a frequency of 0 may represent one to four observed alleles, so raw counts should be checked in gnomAD before a result is relied on.

The alternative to using a program like FIND would be to manually screen gnomAD data for individual variants and apply each of these criteria by hand. This would be an impractical undertaking as it would involve performing multiple operations on hundreds of variants per gene.

**Use of AI tools.** The authors used Claude (Anthropic; Claude Opus 4.5 and 5) to assist with manuscript wording, code debugging and preparation of an earlier pipeline figure. During revision, Claude Opus 5.5 was also used to write the Python scripts for the Fisher’s exact tests, threshold sweep and permutation test (S1–S3 Tables). The scripts reproduced FIND’s output exactly, including all allele frequencies. Script outputs were also validated by recomputing selected p-values independently. The scripts are available as described under Data and Code Availability. The authors reviewed and edited all content and take full responsibility for it; all scientific conceptualization, experimental design and interpretation were performed by the authors.

### Ethics statement

The study used only publicly available aggregate data (gnomAD, ClinVar, All of Us public Data Browser) and required no IRB approval or consent. No individual-level or identifying participant information was accessed.

## Results

We found eleven well-known founder mutations and one that, although not known to be a true founder, was known to be enriched in Ashkenazi Jewish but not in other populations. We also uncovered five variants individually documented as recurrent in specific populations, but none had been previously characterized by systematic cross-population frequency comparison like the analysis FIND performs. That variants documented in specific populations in the literature for years had not previously been compared across populations underscores the analytical gap FIND addresses. Finally, we found three variants not previously documented in specific populations, demonstrating that FIND can also surface previously unreported population-enriched variants (Table 1). These latter two groups are previously unrecognized population-enriched variants, but whether their enrichment reflects a founder effect, selection or another cause is unknown. In categorizing variants by prior literature, population-enriched means a variant whose frequency was compared across populations and found to be higher in one group, as distinct from one reported only as recurrent within a single population without such comparison. FIND’s tenfold enrichment criterion (see Methods) is a separate operational threshold, not the basis for these literature categories. For 13 of the 20 variants, every other group was zeroed, so the tenfold criterion was met automatically; for the other seven, fold ratios ranged from 10.2 to 180 (S1 Table). The population enrichment of all 20 variants was significant by Fisher’s exact test (uncorrected p = 1.3 × 10^-233^ to 1.2 × 10^-5^, all below the Bonferroni-corrected threshold of 2.4 × 10^-5^), including the eight newly reported population-enriched variants (p = 4.1 × 10^-21^ to 4.1 × 10^-7^). In 1,000 permutations, FIND’s criteria returned no variants in 962, one in 37 and two in one, a mean of 0.04 variants per permutation, compared with 26 in the real data (S3 Table).

Loosening the thresholds added variants of mixed quality (S2 Table). A four-allele minimum added one variant that passes Bonferroni correction (MYH7 p.Glu935Lys) and one that fails (BRCA2 c.8633-1299C>G), plus two that our review would remove. Fivefold enrichment added one that narrowly passes (MYH7 p.Arg870Cys) and two that fail (BRCA2 p.Arg3128Ter and p.Leu1768ArgfsTer5). Lowering the minimum allele count further, to three alleles, removed the known *BRCA2* NFE founder from our results, because sporadic low counts in comparison populations were no longer zeroed, raising the comparison frequency and dropping the enrichment ratio below threshold. Raising the minimum to 8 alleles left only 9 of the 12 known and 4 of the 8 newly reported variants. Looser settings increased chance findings: FIND returned at least one variant in 4% of permutations at the defaults, 8% with a four-allele minimum and 21% with fivefold enrichment (S3 Table). We therefore kept the defaults. Users who run FIND from the source code can loosen either threshold and review each additional variant.

The specific variants in each category follow. FIND returned one result for a *Known Population-Enriched* variant in *FLNC* [21–23]. Its enrichment in Ashkenazi Jews was first reported using the IBD Exomes Browser [22], whose Ashkenazi exomes overlap gnomAD, so that comparison is not independent of gnomAD. Independent evidence is its occurrence in three unrelated Ashkenazi Jewish families with arrhythmogenic cardiomyopathy [21], and in another Ashkenazi Jewish family with dilated cardiomyopathy [22]. For *TMEM127* it found a pathogenic variant most common in African or African American populations for which FIND is the *First population-specific report*. The All of Us database showed that the *TMEM127* variant was only present in African ancestry populations, supporting the gnomAD data. Because this variant was observed at a low allele count in gnomAD, its independent appearance in All of Us—again exclusively in individuals of African ancestry—argues that the signal reflects a genuine population-enriched variant rather than a private, single-family variant.

In addition, FIND returned five results for *MYH7*, two of which were pLoF VUS. This is consistent with *MYH7* disease biology: hypertrophic cardiomyopathy is caused predominantly by dominant-negative missense variants rather than haploinsufficiency, so most truncating variants are not pathogenic. In gnomAD, 96 of 101 classified LoF variants in *MYH7* are VUS and only 5 are classified as pathogenic. We therefore did not pursue our two pLoF variant hits further. Of the remaining three hits, neither c.2606G>A (enriched in African American) nor c.1491G>T (enriched in Ashkenazi Jewish) had been previously reported as a founder or as population-enriched; c.2606G>A had, however, been reported as recurrent in a Tuscan cohort [25] without comparison to other populations (FIND provides the *First cross-population comparison)*; whereas c.1491G>T had no prior report making it a *First population-specific report*. All of Us corroborates that one of these findings (c.2606G>A) is enriched in the African American population. The remaining pathogenic *MYH7* variant was previously identified as a Finnish *Known Founder* [24].

FIND returned five results for *BRCA1*. Two variants were Ashkenazi Jewish *Known Founder* mutations, one was an East Asian *Known Founder* [26]. Another one was previously reported in Japanese patients [27] but enrichment relative to other populations was not examined. FIND provides the *First cross-population comparison* for this variant. The remaining variant was excluded because it failed gnomAD’s allele-specific Variant Quality Score Recalibration (AS-VQSR) filter, indicating a likely artifact. Because *BRCA1* was analyzed only after the thresholds were set, FIND’s recovery of three *BRCA1* founders listed in a review [26] is an independent test of FIND.

Finally, FIND returned 14 results for *BRCA2*. Three were characterized as VUS. In contrast with *MYH7,* loss of function is the established *BRCA2* disease mechanism and almost all loss of function *BRCA2* variants in gnomAD are pathogenic. The three VUS pLoF variants obtained here are exceptions because they fall near the 3′ end of the gene, where premature stop codons escape nonsense-mediated decay and yield a near-full-length protein, and are therefore not expected to actually cause loss of function. Therefore, we did not pursue these variants further. More broadly, these examples illustrate that loss-of-function status alone does not establish pathogenicity: its disease relevance depends on the gene’s mechanism and on the mutation’s position in the gene, which is why FIND flags pLoF variants as candidates for evaluation. Of the remaining eleven, seven are *Known Founders*: an Ashkenazi Jewish founder c.5946del [34], an Admixed American founder c.3922G>T [35], an East Asian founder c.5576_5579del [36], a Non-Finnish European founder c.6275_6276del [37], and two Finnish founders, c.7480C>T and c.9118-2A>G [29]. Included in the seven, FIND recovered the known Icelandic founder (c.771_775del) which gnomAD surfaces in the Finnish group rather than as Icelandic, since gnomAD has no Icelandic ancestry group. This variant also segregates in Finnish breast cancer families [29] consistent with its enrichment in that group.

Among the remaining four *BRCA2* results, three were previously documented as recurrent in specific populations but had not been compared across populations (labelled *First cross-population comparison)*. One was described in three Japanese patients [38], two were common in Hispanic populations (c.3264dup [30, 31] and c.9235del [32, 33]). The fourth variant, p.Cys3233TrpfsTer15, was revealed to be enriched in Admixed Americans by FIND but had not been previously reported in a specific population (*First population-specific report*). The variants FIND reports as population-enriched in Admixed American groups are likewise population-enriched in the All of Us database as seen in the table. Finding that a variant is population-enriched in two separate unrelated data bases is strong support that the enrichment is genuine: because the two databases differ in recruitment and geographic representation, independent replication makes a database-specific artifact an unlikely explanation.

## Discussion

Statistical analyses support FIND’s results. Every variant in Table 1 was significant by Fisher’s exact test on raw allele counts, even after correction for multiple testing, and a permutation test showed that FIND’s criteria rarely return a variant by chance. Varying the allele-count and enrichment thresholds showed that the default settings retain all known variants, while looser settings add variants of mixed quality. Apart from FLNC c.3791-1G>C (see Results), every Known Founder and Known Population-Enriched label in Table 1 rests on population-specific clinical or cohort studies independent of gnomAD, so FIND’s recovery of these variants shows that its screen detects enrichment documented in other data.

To reiterate, FIND simply highlights variants that are enriched in a gnomAD-specific ancestry population without establishing that they are true founder mutations. To do that, these variants would need to undergo haplotype analysis [2]. A variant can be common in one group for more than one reason. It may descend from a single ancestor, as in a founder population. It may have been kept in that group while other groups lost it. Or it may have been favored by selection. Allele frequency alone cannot tell these apart. FIND relies on allele frequency data.

Ancestry groupings in gnomAD are broad categories that imperfectly capture genetic diversity within populations and some populations in gnomAD have had many more sequences recorded than others. This matters because founder mutations arise in small communities, while most gnomAD groups cover whole continents. A variant that is common in one community can look uncommon once it is averaged over an entire group. In addition, FIND uses what ClinVar reports for pathogenic and likely pathogenic classifications, which are not always consensus calls. As gnomAD expands in size and refines its ancestry definitions and as the pathogenicity classifications of ClinVar improve, the accuracy of FIND’s outputs will improve accordingly. Indeed, with the growing availability of large-scale whole-exome and whole-genome datasets ancestry groups can increasingly be resolved at a finer scale, reducing ambiguities that arise when admixed individuals are assigned to broad, discrete groups.

Some limitations of the FIND program are illustrated by the *BRCA2* c.7480C>T variant. This variant has the unusual characteristic of being previously identified as a founder mutation in two different populations, Finnish [29] and Korean [39]. The appearance of *BRCA2* c.7480C>T in two unrelated founder populations is consistent with recurrent mutation at a CpG dinucleotide hotspot [40]. The low frequency of Koreans included in the gnomAD database [41] prevented its identification as an East Asian founder; moreover, the inclusion of Koreans into the East Asian population masks Korean-specific variants. Furthermore, FIND’s requirement that an enriched population frequency be ten times higher than in any other population prevents the identification of a variant that is a founder in two different populations. It was detected only as a Finnish founder because its East Asian count of two was zeroed under our threshold. Variants that arose before populations diverged are similarly missed for failing the tenfold-enrichment criterion. Thus, FIND has difficulties detecting population-enriched variants from populations with little data in gnomAD, populations that are inaccurately classified in gnomAD, and variants that are enriched in multiple populations.

To explore FIND’s sensitivity, we compared its output to a published review of *BRCA1/2* founder mutations [26]. For *BRCA1,* FIND recovered the two Ashkenazi Jewish founders and one East Asian founder listed in the review. The Finnish founder listed is now categorized as a VUS and was very rare in gnomAD. For *BRCA2,* FIND recovered one Ashkenazi Jewish, one Finnish, one East Asian, and one non-Finnish European founder listed in the review as well as the listed known Icelandic founder (c.771_775del alias 999del5), which gnomAD surfaces in the Finnish group as described in **Results**. Other *BRCA1/2* founders listed in the review [26] were either missing or very rare in gnomAD. In one case a Swedish variant with 11 non-Finnish European copies in gnomAD had too low a frequency to pass our screen because of being drowned out by the other populations in the non-Finnish European group. Table 1 includes two additional known *BRCA2* founders not listed in the review [26] but described in more recent population-specific literature. This pattern reinforces FIND’s central limitation, dependence on gnomAD data. When the minimum allele count was raised, newly reported variants dropped out faster than known variants (S2 Table), as expected for variants rare enough to have escaped earlier notice.

Beyond research utility, knowledge of population-enriched pathogenic variants has potential clinical benefits. Despite tremendous recent advances in understanding connections between mutations and human disease, real-world implementation lags behind expert clinical guidelines. For example, currently only 8% of cardiomyopathy patients that guidelines recommend get genetic testing are actually prescribed testing by their doctors, and only 1% proceed to get tested; furthermore, when a pathogenic variant is uncovered only about 33% proceed to cascade testing [42]. We propose that knowing that a pathogenic variant is enriched in a patient’s ancestral population will help motivate physicians to order testing in the first place and will motivate patients and families to pursue cascade testing once a variant is found. This is illustrated by recent efforts to raise awareness of a *PLN* founder mutation among individuals of Dutch ancestry in West Michigan, where the mutation is thought to be particularly prevalent [43].

A recent example where population data raised awareness is a nonsense variant in CD36 (rs3211938, p.Tyr325Ter), carried by about 17% of African ancestry individuals but rare in other groups [44]. Homozygotes have a three-to five-fold higher risk of dilated cardiomyopathy and heterozygotes a modestly higher risk [44, 45]. FIND returns it as enriched in the African/African American group (allele frequency 0.091) even though ClinVar classifies it as uncertain significance or benign, because it is a predicted loss-of-function variant. Until now it had not been included in standard cardiomyopathy testing panels. Cleveland Clinic plans to rectify that, because it is now clear that the variant is a major source of dilated cardiomyopathy risk in African/African Americans [45, 46].

FIND may also help address disparities in variant identification across populations. Our identification of population-enriched variants in *TMEM127, MYH7,* an*d BRCA2*, in African/African American or Admixed American populations and corroborated in All of Us (whose oversampling of historically under-represented populations makes such independent corroboration possible precisely where gnomAD sample sizes are smallest), demonstrates how FIND can identify clinically relevant population-enriched variants in understudied groups. FIND replaces an otherwise manual process and offers a principled approach to uncovering population-enriched pathogenic variants, whose utility will grow as gnomAD and ClinVar continue to improve.

## Acknowledgments

We thank Drs. Gai Elhanan and Brian Shirts and the anonymous PLOS ONE reviewer for helpful suggestions and comments on the manuscript. We gratefully acknowledge All of Us participants for their contributions, without whom this research would not have been possible. We also thank the National Institutes of Health’s All of Us Research Program for making available the aggregate data examined in this study.

## Author Contributions

A.L.H. and S.W.L. contributed to conceptualization, formal analysis, investigation, methodology, validation, and writing. A.Z.L. designed and wrote the FIND software, tested various genes during development, debugged and refined the implementation, and contributed technical inputs used in figure preparation. A.L.H. performed extensive testing with five ACMG-actionable genes, and led the data interpretation, manuscript drafting, and designed the figures. S.W.L. selected the genes used for software development and systematic testing and provided supervision. S.W.L. performed the statistical analyses (S1–S3 Tables). All authors reviewed and approved the final manuscript.

## Data and Code Availability

The FIND source code is freely available under an MIT license on GitHub (https://github.com/aacoder105/FIND) and permanently archived at Zenodo (https://doi.org/10.5281/zenodo.20499975). A web interface is hosted on Render (https://ethnic-variant-mutation-finder.onrender.com/). Users can also deploy FIND locally. FIND queries gnomAD v4.1 (https://gnomad.broadinstitute.org/), which is freely available; pathogenicity classifications are the ClinVar annotations provided within gnomAD, from ClinVar (https://www.ncbi.nlm.nih.gov/clinvar/), also freely available. All of Us aggregate allele-frequency data used are available through the public Data Browser at https://databrowser.researchallofus.org/ (CDR v8 release). The analysis scripts and gnomAD data files used for the analyses in S1–S3 Tables are available at https://github.com/aacoder105/FIND (folder FIND_revision_analysis_v2/FIND_revision_analysis) and archived at Zenodo (https://doi.org/10.5281/zenodo.20499975).

## Supporting Information

S1 Table. Raw gnomAD allele counts, fold enrichment and Fisher’s exact tests for the 20 variants in Table 1.

S2 Table. Effect of FIND thresholds on results for the five test genes, and additional variants returned at relaxed settings.

S3 Table. Permutation test of FIND results expected by chance at default and relaxed settings.

## References

1. Gombault C, Grenet G, Segurel L, Duret L, Gueyffier F, Cathebras P, et al. Population designations in biomedical research: Limitations and perspectives. HLA. 2023;101(1):3–15. Epub 20221105. doi: 10.1111/tan.14852. PubMed PMID: 36258305; PubMed Central PMCID: PMC10099491.

2. Neuhausen SL, Mazoyer S, Friedman L, Stratton M, Offit K, Caligo A, et al. Haplotype and phenotype analysis of six recurrent BRCA1 mutations in 61 families: results of an international study. Am J Hum Genet. 1996;58(2):271–80. PubMed PMID: 8571953; PubMed Central PMCID: PMC1914544.

3. Jain A, Sharma D, Bajaj A, Gupta V, Scaria V. Founder variants and population genomes-Toward precision medicine. Adv Genet. 2021;107:121–52. Epub 20210218. doi: 10.1016/bs.adgen.2020.11.004. PubMed PMID: 33641745.

4. Tennen RI, Laskey SB, Koelsch BL, McIntyre MH, Tung JY. Identifying Ashkenazi Jewish BRCA1/2 founder variants in individuals who do not self-report Jewish ancestry. Sci Rep. 2020;10(1):7669. Epub 20200506. doi: 10.1038/s41598-020-63466-x. PubMed PMID: 32376921; PubMed Central PMCID: PMC7203114.

5. Behl S, Hamel N, de Ladurantaye M, Lepage S, Lapointe R, Mes-Masson AM, et al. Founder BRCA1/BRCA2/PALB2 pathogenic variants in French-Canadian breast cancer cases and controls. Sci Rep. 2020;10(1):6491. Epub 20200416. doi: 10.1038/s41598-020-63100-w. PubMed PMID: 32300229; PubMed Central PMCID: PMC7162921.

6. Kandolin M, Poyhonen M, Jakkula E. Estimation of carrier frequencies utilizing the gnomAD database for ACMG recommended carrier screening and Finnish disease heritage conditions in non-Finnish European, Finnish, and Ashkenazi Jewish populations. Am J Med Genet A. 2024;194(7):e63588. Epub 20240308. doi: 10.1002/ajmg.a.63588. PubMed PMID: 38459613.

7. Fatumo S, Chikowore T, Choudhury A, Ayub M, Martin AR, Kuchenbaecker K. A roadmap to increase diversity in genomic studies. Nat Med. 2022;28(2):243–50. Epub 20220210. doi: 10.1038/s41591-021-01672-4. PubMed PMID: 35145307; PubMed Central PMCID: PMC7614889.

8. Abe TA, Lancaster MC, Roden DM. Association of Common Ancestry-Enriched Variants With Cardiomyopathy and Arrhythmias. Circulation. 2026;153(24):1915–27. Epub 20260527. doi: 10.1161/CIRCULATIONAHA.126.080245. PubMed PMID: 42200287.

9. Foulkes WD, Knoppers BM, Turnbull C. Population genetic testing for cancer susceptibility: founder mutations to genomes. Nat Rev Clin Oncol. 2016;13(1):41–54. Epub 20151020. doi: 10.1038/nrclinonc.2015.173. PubMed PMID: 26483301.

10. Kore P, Wilson MW, Tiao G, Chao K, Darnowsky PW, Watts NA, et al. Improved allele frequencies in gnomAD through local ancestry inference. Nat Commun. 2025;16(1):8734. Epub 20251006. doi: 10.1038/s41467-025-63340-2. PubMed PMID: 41053080; PubMed Central PMCID: PMC12500861.

11. Guez J, Goodrich JK, Moldovan MA, Chao KR, Kar P, Panchal R, et al. Integrating 730,947 exome sequences with clinical literature improves gene discovery. medRxiv. 2026. Epub 20260325. doi: 10.64898/2026.03.23.26349081. PubMed PMID: 41929314; PubMed Central PMCID: PMC13042128.

12. Landrum MJ, Lee JM, Benson M, Brown G, Chao C, Chitipiralla S, et al. ClinVar: public archive of interpretations of clinically relevant variants. Nucleic Acids Res. 2016;44(D1):D862–8. Epub 20151117. doi: 10.1093/nar/gkv1222. PubMed PMID: 26582918; PubMed Central PMCID: PMC4702865.

13. Karczewski KJ, Francioli LC, Tiao G, Cummings BB, Alfoldi J, Wang Q, et al. The mutational constraint spectrum quantified from variation in 141,456 humans. Nature. 2020;581(7809):434–43. Epub 20200527. doi: 10.1038/s41586-020-2308-7. PubMed PMID: 32461654; PubMed Central PMCID: PMC7334197.

14. Chen S, Francioli LC, Goodrich JK, Collins RL, Kanai M, Wang Q, et al. A genomic mutational constraint map using variation in 76,156 human genomes. Nature. 2024;625(7993):92–100. Epub 20231206. doi: 10.1038/s41586-023-06045-0. PubMed PMID: 38057664; PubMed Central PMCID: PMC11629659.

15. Hatchell KE, Poll SR, Russell EM, Williams TJ, Ellsworth RE, Facio FM, et al. Experience using conventional compared to ancestry-based population descriptors in clinical genomics laboratories. Am J Hum Genet. 2025;112(3):481–91. Epub 20250129. doi: 10.1016/j.ajhg.2025.01.008. PubMed PMID: 39884281; PubMed Central PMCID: PMC11947177.

16. Horowitz AL, Liebman AZ, Liebman SW. FIND: a software tool for identifying population-enriched pathogenic variants in gnomAD. bioRxiv [Preprint] 2026. Version 1, posted 2026.06.05. doi:10.64898/2026.06.05.730273.

17. Amberger JS, Bocchini CA, Scott AF, Hamosh A. OMIM.org: leveraging knowledge across phenotype-gene relationships. Nucleic Acids Res. 2019;47(D1):D1038–D43. doi: 10.1093/nar/gky1151. PubMed PMID: 30445645; PubMed Central PMCID: PMC6323937.

18. All of Us Research Program Genomics Investigators. Genomic data in the All of Us Research Program. Nature. 2024;627(8003):340–6. Epub 20240219. doi: 10.1038/s41586-023-06957-x. PubMed PMID: 38374255; PubMed Central PMCID: PMC10937371.

19. All of Us Research Program Genomics Investigators. All of Us Genomic Quality Report [Internet]. Controlled Tier CDR v8; released 2025 Feb 3 [cited 2026 Jun 21]. 2025. Available from: https://support.researchallofus.org/hc/en-us/articles/29390274413716-All-of-Us-Genomic-Quality-Report.

20. Lee K, Abul-Husn NS, Amendola LM, Brothers KB, Chung WK, Gollob MH, et al. ACMG SF v3.3 list for reporting of secondary findings in clinical exome and genome sequencing: A policy statement of the American College of Medical Genetics and Genomics (ACMG). Genet Med. 2025;27(8):101454. Epub 20250623. doi: 10.1016/j.gim.2025.101454. PubMed PMID: 40568962; PubMed Central PMCID: PMC12318660.

21. Oz S, Yonath H, Visochyk L, Ofek E, Landa N, Reznik-Wolf H, et al. Reduction in Filamin C transcript is associated with arrhythmogenic cardiomyopathy in Ashkenazi Jews. Int J Cardiol. 2020;317:133–8. Epub 20200405. doi: 10.1016/j.ijcard.2020.04.005. PubMed PMID: 32532510.

22. Liebman SW, Palaganas H, Kobany H. A founder mutation in FLNC is likely a major cause of idiopathic dilated cardiomyopathy in Ashkenazi Jews. Int J Cardiol. 2021;323:124. Epub 20200815. doi: 10.1016/j.ijcard.2020.08.052. PubMed PMID: 32805325.

23. Deo RC, Musso G, Tasan M, Tang P, Poon A, Yuan C, et al. Prioritizing causal disease genes using unbiased genomic features. Genome Biol. 2014;15(12):534. Epub 20141203. doi: 10.1186/s13059-014-0534-8. PubMed PMID: 25633252; PubMed Central PMCID: PMC4279789.

24. Jaaskelainen P, Helio T, Aalto-Setala K, Kaartinen M, Ilveskoski E, Hamalainen L, et al. A new common mutation in the cardiac beta-myosin heavy chain gene in Finnish patients with hypertrophic cardiomyopathy. Ann Med. 2014;46(6):424–9. Epub 20140603. doi: 10.3109/07853890.2014.912834. PubMed PMID: 24888384.

25. Mazzarotto F, Girolami F, Boschi B, Barlocco F, Tomberli A, Baldini K, et al. Defining the diagnostic effectiveness of genes for inclusion in panels: the experience of two decades of genetic testing for hypertrophic cardiomyopathy at a single center. Genet Med. 2019;21(2):284–92. Epub 20180606. doi: 10.1038/s41436-018-0046-0. PubMed PMID: 29875424; PubMed Central PMCID: PMC6752309.

26. Ferla R, Calo V, Cascio S, Rinaldi G, Badalamenti G, Carreca I, et al. Founder mutations in BRCA1 and BRCA2 genes. Ann Oncol. 2007;18 Suppl 6:vi93–8. doi: 10.1093/annonc/mdm234. PubMed PMID: 17591843.

27. Yoshida R, Watanabe C, Yokoyama S, Inuzuka M, Yotsumoto J, Arai M, et al. Analysis of clinical characteristics of breast cancer patients with the Japanese founder mutation BRCA1 L63X. Oncotarget. 2019;10(35):3276–84. Epub 20190514. doi: 10.18632/oncotarget.26852. PubMed PMID: 31143373; PubMed Central PMCID: PMC6524931.

28. Thorlacius S, Sigurdsson S, Bjarnadottir H, Olafsdottir G, Jonasson JG, Tryggvadottir L, et al. Study of a single BRCA2 mutation with high carrier frequency in a small population. Am J Hum Genet. 1997;60(5):1079–84. PubMed PMID: 9150155; PubMed Central PMCID: PMC1712443.

29. Sarantaus L, Huusko P, Eerola H, Launonen V, Vehmanen P, Rapakko K, et al. Multiple founder effects and geographical clustering of BRCA1 and BRCA2 families in Finland. Eur J Hum Genet. 2000;8(10):757–63. doi: 10.1038/sj.ejhg.5200529. PubMed PMID: 11039575.

30. Jara L, Morales S, de Mayo T, Gonzalez-Hormazabal P, Carrasco V, Godoy R. Mutations in BRCA1, BRCA2 and other breast and ovarian cancer susceptibility genes in Central and South American populations. Biol Res. 2017;50(1):35. Epub 20171006. doi: 10.1186/s40659-017-0139-2. PubMed PMID: 28985766; PubMed Central PMCID: PMC6389095.

31. Rebbeck TR, Friebel TM, Friedman E, Hamann U, Huo D, Kwong A, et al. Mutational spectrum in a worldwide study of 29,700 families with BRCA1 or BRCA2 mutations. Hum Mutat. 2018;39(5):593–620. Epub 20180312. doi: 10.1002/humu.23406. PubMed PMID: 29446198; PubMed Central PMCID: PMC5903938.

32. Ferreyra Y, Rosas G, Cock-Rada AM, Araujo J, Bravo L, Doimi F, et al. Landscape of germline BRCA1/BRCA2 variants in breast and ovarian cancer in Peru. Front Oncol. 2023;13:1227864. Epub 20230817. doi: 10.3389/fonc.2023.1227864. PubMed PMID: 37664050; PubMed Central PMCID: PMC10470619.

33. Villarreal-Garza C, Alvarez-Gomez RM, Perez-Plasencia C, Herrera LA, Herzog J, Castillo D, et al. Significant clinical impact of recurrent BRCA1 and BRCA2 mutations in Mexico. Cancer. 2015;121(3):372–8. Epub 20140918. doi: 10.1002/cncr.29058. PubMed PMID: 25236687; PubMed Central PMCID: PMC4304938.

34. Abeliovich D, Kaduri L, Lerer I, Weinberg N, Amir G, Sagi M, et al. The founder mutations 185delAG and 5382insC in BRCA1 and 6174delT in BRCA2 appear in 60% of ovarian cancer and 30% of early-onset breast cancer patients among Ashkenazi women. Am J Hum Genet. 1997;60(3):505–14. PubMed PMID: 9042909; PubMed Central PMCID: PMC1712523.

35. Abul-Husn NS, Soper ER, Odgis JA, Cullina S, Bobo D, Moscati A, et al. Exome sequencing reveals a high prevalence of BRCA1 and BRCA2 founder variants in a diverse population-based biobank. Genome Med. 2019;12(1):2. Epub 20191231. doi: 10.1186/s13073-019-0691-1. PubMed PMID: 31892343; PubMed Central PMCID: PMC6938627.

36. Nakamura H, Mizumoto S, Tanino H, Niwa Y, Ogino M, Sakoda Y, et al. High Frequency of BRCA2 c.5576_5579del Carriers in Kakogawa, Japan. Cancer Diagn Progn. 2024;4(3):309–14. Epub 20240503. doi: 10.21873/cdp.10325. PubMed PMID: 38707742; PubMed Central PMCID: PMC11062150.

37. Scottish/Northern Irish BRCA1/BRCA2 Consortium. BRCA1 and BRCA2 mutations in Scotland and Northern Ireland. Br J Cancer. 2003;88(8):1256–62. doi: 10.1038/sj.bjc.6600840. PubMed PMID: 12698193; PubMed Central PMCID: PMC2747571.

38. Arai M, Yokoyama S, Watanabe C, Yoshida R, Kita M, Okawa M, et al. Genetic and clinical characteristics in Japanese hereditary breast and ovarian cancer: first report after establishment of HBOC registration system in Japan. J Hum Genet. 2018;63(4):447–57. Epub 20171108. doi: 10.1038/s10038-017-0355-1. PubMed PMID: 29176636; PubMed Central PMCID: PMC8716335.

39. Seong MW, Cho S, Noh DY, Han W, Kim SW, Park CM, et al. Comprehensive mutational analysis of BRCA1/BRCA2 for Korean breast cancer patients: evidence of a founder mutation. Clin Genet. 2009;76(2):152–60. Epub 20090728. doi: 10.1111/j.1399-0004.2009.01202.x. PubMed PMID: 19656164.

40. Cooper DN, Krawczak M. The mutational spectrum of single base-pair substitutions causing human genetic disease: patterns and predictions. Hum Genet. 1990;85(1):55–74. doi: 10.1007/BF00276326. PubMed PMID: 2192981.

41. Lee J, Lee J, Jeon S, Lee J, Jang I, Yang JO, et al. A database of 5305 healthy Korean individuals reveals genetic and clinical implications for an East Asian population. Exp Mol Med. 2022;54(11):1862–71. Epub 20221102. doi: 10.1038/s12276-022-00871-4. PubMed PMID: 36323850; PubMed Central PMCID: PMC9628380.

42. Genome Medical. Voice of the Patient 2026 [June 21, 2026]. Available from: https://www.genomemedical.com/resources/voice-of-patient/cardiomyopathy/.

43. WZZM13. Do you have Dutch ancestry? A rare gene mutation could affect your heart 2026 [June 21, 2026]. Available from: https://www.wzzm13.com/article/news/local/do-you-have-dutch-ancestry-rare-gene-mutation-could-affect-heart/69-7a8010ee-1950-4e39-b554-31ad6ea01f04.

44. Huffman JE, Gaziano L, Al Sayed ZR, Judy RL, Raffield LM, Biddinger KJ, et al. An African ancestry-specific nonsense variant in CD36 is associated with a higher risk of dilated cardiomyopathy. Nat Genet. 2025;57(11):2682–90. Epub 20251031. doi: 10.1038/s41588-025-02372-2. PubMed PMID: 41174181; PubMed Central PMCID: PMC12597818.

45. Challa SP, Biddinger KJ, Abramowitz SA, Zheng A, Mead JO, Judy RL, et al. A Common CD36 Variant and the Genetic Landscape of Dilated Cardiomyopathy in Individuals of African Ancestry. JAMA. 2026. Epub 20261019. doi: 10.1001/jama.2026.18449. PubMed PMID: 42895716.

46. Cleveland Clinic. Cleveland Clinic-led study establishes major genetic contributor to heart failure in people of African ancestry 2026 [October 20, 2026]. Available from: https://newsroom.clevelandclinic.org/2026/10/19/cleveland-clinic-led-study-establishes-major-genetic-contributor-to-heart-failure-in-people-of-african-ancestry.

